# The molecular architecture of the aquaglyceroporin gating system in *Saccharomyces cerevisiae*

**DOI:** 10.1101/2025.10.24.684117

**Authors:** Mikael Andersson, Linde Viaene, Adeleine Marie Albalate, Mark C Leake, Adam Wollman, Sviatlana Shashkova

**Affiliations:** Department of Chemistry and Molecular Biology, University of Gothenburg, Gothenburg, Sweden; Department of Physics, University of Gothenburg, Sweden; School of Physics, Engineering and Technology, University of York, York, UK; Department of Biology, University of York, York; Biosciences Institute, Newcastle University, Newcastle-upon-Tyne, UK

## Abstract

Aquaglyceroporins have been extensively studied in multiple systems. However, the molecular architecture and hence the precise mechanisms of action remain unknow. The yeast aquaglyceroporin Fps1 plays the main role in controlling cellular turgor. It is regulated by the two Rgc proteins which activate the Fps1 channel. To date, the molecular architecture and dynamics of the Fps1-Rgc gating system is unknown. We employed dual-coloured super-resolution millisecond fluorescence microscopy with a single-molecule sensitivity, to track Fps1 and Rgc2 in real time. Unlike previously believed, our data shows that Rgc2 mainly operates as a homodimer. At the same time, Rgc1, the seemingly redundant regulatory ortholog of Rgc2, has a clear effect on the architecture, colocalisation behaviour and molecular dynamics of the Fps1 and Rgc2 together. Moreover, Rgc1 seems to promote Rgc2 stability in response to hydrogen peroxide stress. Together, our results provide novel insights into the mechanism behind aquaglyceroporins regulation.

**Importance:** The components of the Fps1 aquaglyceroporin gating system is conserved in Fungi and present in both baker’s yeast *Saccharomyces cerevisiae* as well as common pathogenic included in the WHO fungal priority pathogens list: *Candida albicans*, *Candida glabrata, Aspergillus fumigatus, Pichia kudriavzevii*. For instance, in *C. glabrata*, the orthologs of both Fps1 and Rgc2 are involved in key pathogenic processes related to acetate tolerance, phagocytosis by macrophages, resistance to antifungal agents and overall drug uptake and efflux. As the Fps1 gating system is not present in metazoans, it constitutes an attractive drug target for antifungal agents. Thus, understanding the architecture and the finer mechanisms of the Fps1 pore regulation is a stepping stone in development of effective antifungal treatment and targeted antifungal drugs.

## Introduction

Around half of the cell membrane content is composed of proteins with critical functions. This enables cells to communicate with their environment and maintain an intracellular milieu permissive to life. One of the most common types of membrane proteins which works directly at the interface between the intracellular and extracellular environment are aquaglyceroporins (1). This subtype of the larger aquaporin family facilitates the passive diffusion of small molecules including uncharged solutes in yeast and other organisms (2–5). Fps1 is a tightly regulated yeast aquaglyceroporin that can transport glycerol and a range of other biologically relevant small molecules, such as acetate and toxic metalloids (6–12). Rgc1 and Rgc2 are two mutually redundant paralogous proteins integral to the Fps1 channel activity. Rgc1 and Rgc2 are encoded by two paralogous genes, and either of those genes are sufficient to enable functional Fps1 regulation (13–15). In addition, Rgc proteins have been shown to bind to each other to functionally control Fps1 as homo- or heterodimers (16). This binding to Fps1 along with accompanying Fps1 phosphorylation events enables the active state of the channel (Fig. 1) (14, 17, 18).

**Figure 1.**
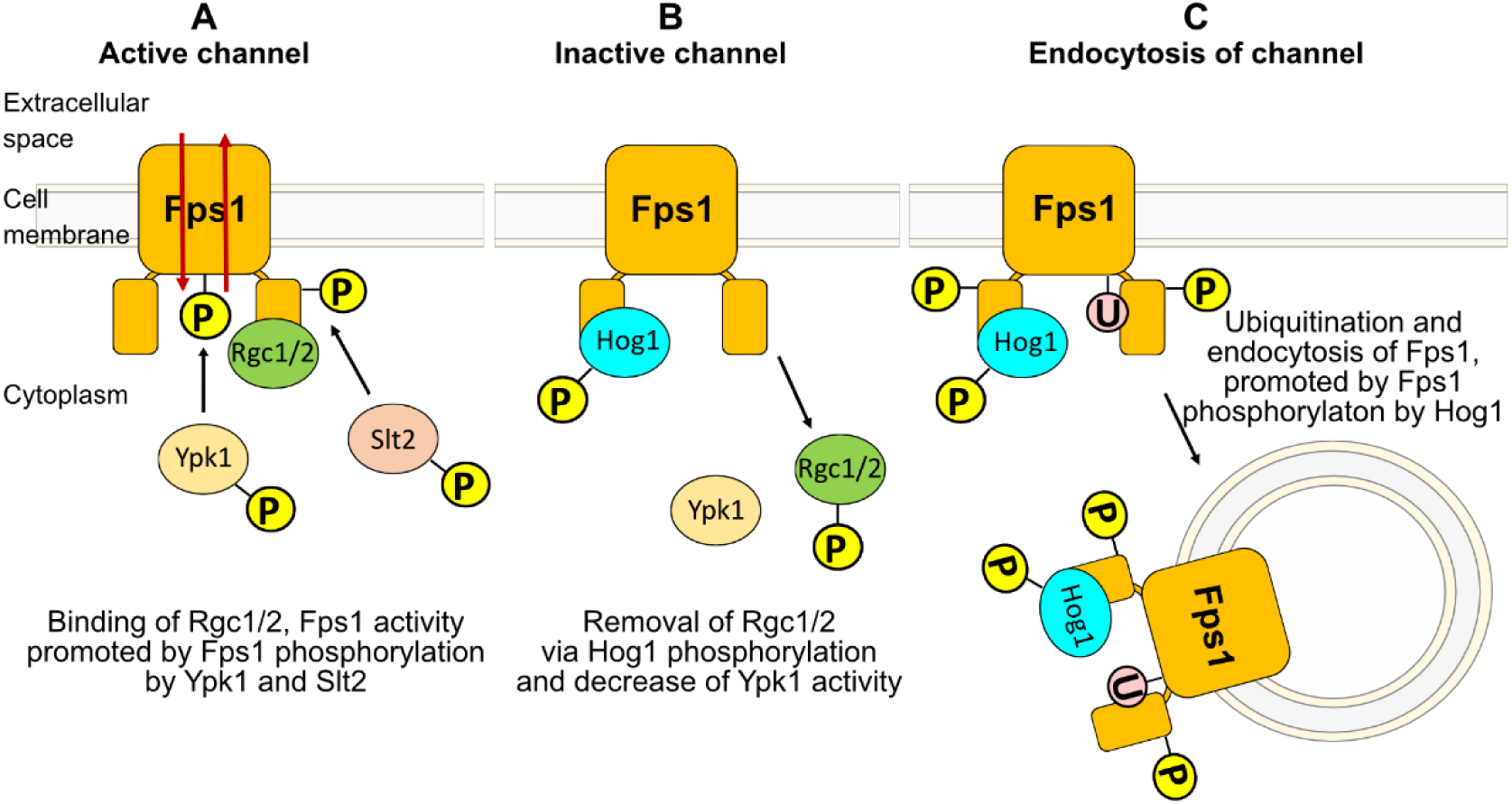
Schematic representation of the Fps1 regulatory states. As the cell needs to control its turgor, it is thought that Fps1 needs to be continuously regulated throughout the cell cycle to cope with osmotic challenges and recycle protein. **A**. The main open active state of Fps1, requires binding of Rgc1 and Rgc2 proteins as well as phosphorylation by the Ypk1 and Slt2 kinases. This typically happens in response to hypoosmotic stress, when the cell needs to relieve intracellular turgor. **B.** For the closed inactive state, the Hog1 kinase binds and phosphorylates Fps1 at the N-terminus and evicts Rgc2 via hyperphosphorylation. At the same time, Ypk1 effect on Fps1 decreases. This is the response when intracellular turgor needs to be increased, e.g., upon hyperosmotic stress or when the channel needs to be closed to prevent the influx of toxic compounds through Fps1. **C**. Fps1 endocytosis is known to involve Hog1-dependent specific N- and C-terminal Fps1 phosphorylation followed by ubiquitination. Fps1 endocytosis has been linked to oxidative stress induced by acetate.

Fps1 regulation is closely associated with the HOG pathway, where the Hog1 protein kinase plays the central role(19). Hog1 is a Mitogen-Activated Protein Kinase (MAPK) that can bind Fps1 and downregulate its’ activity by phosphorylation as well as phosphorylating Rgc1 and Rgc2 proteins leading to their dissociation from Fps1 (14). This cascade of events is thought to close the channel (14), e.g., upon hyperosmotic stress where the cell needs to retain intracellular glycerol to regain normal turgor or arsenite stress where Fps1 needs to be closed to prevent influx of toxic arsenite (14, 15). Hog1 phosphorylation of Fps1 is also known to induce the endocytosis of the channel (9). Fps1 is posited to have a low level of continual activity during normal growth (6, 20). This low-level activity has been associated with the need for glycerol release to prevent cell wall stress due to glycerol being continuously generated via cellular metabolism (6, 20, 21). Fps1 also has a constant turnover (15). This along with the low-level continual activity suggest that Fps1 naturally cycles through various states of regulation under normal growth (Fig. 1).

The components of the Fps1-Rgc1/Rgc2 transport system are also present in common pathogenic fungi, including *Candida albicans*, *Candida glabrata, Aspergillus fumigatus, Pichia kudriavzevii* – all on the World Health Organisation (WHO) fungal priority pathogens list (22). The functional gated pore system with Fps1 and Rgc1/Rgc2 is largely conserved between *C. glabrata* and the better studied *Saccharomyces cerevisiae*. In *C. glabrata*, the orthologs of both Fps1 and Rgc2 are involved in key pathogenic processes related to acetate tolerance (23, 24), phagocytosis by macrophages (23), resistance to antifungal agents targeting the cell wall (25) and overall drug uptake and efflux (26). The Fps1-Rgc1/Rgc2 system is not present in metazoans (16), hence, it constitutes an attractive drug target for antifungal agents. Thus, understanding the architecture and the finer mechanisms of the Fps1-Rgc1/Rgc2 gated pore regulation is important to the development of effective antifungal treatment and targeted antifungal drugs.

The genetic and physical interactions between Fps1 and Rgc1/Rgc2 have previously been studied *in vitro* and *in vivo* on a population level (14, 15, 17, 27, 28). This has yielded information regarding the critical importance, and apparent redundancy, of Rgc1/Rgc2 in regulating Fps1 activity during osmotic and metalloid stresses through their binding to, or removal from, Fps1. However, more detailed focus on molecules within individual cells enables detection of cell-to-cell variability and molecular subpopulations which may perform different functions and hence play a crucial role in cellular adaptation and survival in different conditions (29). Our recent work on the molecular characteristics of Fps1 revealed a previously unseen intracellular pool of the protein which exists as mobile and immobile fractions. Moreover, we showed that Fps1 resides in multi-tetrameric assemblies and that subjection to H_2_O_2_ stress causes Fps1 degradation (30).

As the presence of Rgc proteins on Fps1 is correlated with the opening of the channel and hence Fps1 activity, thus, observing colocalisation between these proteins serves as a proxy for observing Fps1 opening *in vivo*. The molecular architecture of Rgc proteins within the active Fps1 gating system in cells is also unknown. Therefore, we created a set of dual-coloured strains (Supplementary Fig.1) and employed single-molecule super-resolution Slimfield fluorescence microscopy (31) to determine the architecture, biophysical characteristics and colocalisation dynamics between Fps1 and Rgc2 in the budding yeast *S. cerevisiae.* We visualised the molecular architecture of fluorescently labelled proteins in response to genetic and osmotic perturbation to obtain key functional insights into the behaviour of Fps1 and Rgc2 separately and as a complex. Our data shows that Rgc2 mainly operates as a homodimer. At the same time, Rgc1, the seemingly redundant regulatory ortholog of Rgc2, have a clear effect on the architecture, colocalisation behaviour and molecular dynamics of the Fps1 and Rgc2. Moreover, Rgc1 seems to promote Rgc2 stability in response to hydrogen peroxide stress.

## Results

### Fps1 and Rgc2 colocalise both intracellularly and on the cell membrane

We employed dual-coloured single-molecule Slimfield fluorescence microscopy to visualise individual fluorescent foci and thereby determine the architecture and localisations of Fps1 and Rgc2 simultaneously. Slimfield microscopy utilises a slim laser excitation field to capture single-molecule fluorescence at high time resolution and has been widely used in bacteria (32–35) and yeast (36–39). We used a dual-labelled yeast strain containing endogenously expressed Fps1-mGFP and Rgc2-mCherry protein constructs (Fig. 2A). Similarly to the approach in our previous study (30), any GFP or mCherry foci found at the cell boundaries identified from the brightfield, we further refer to as “membrane”, while the spots found in the rest of the cell as “intracellular”. We observed between 1-17 Fps1 foci per cell, with a mean of 8 (Supplementary Table 1). Simultaneous imaging of Rgc2-mCherry in these cells indicated between 2-25 foci per cell, with a mean of 10 (Supplementary Table 1).

**Figure 2.**
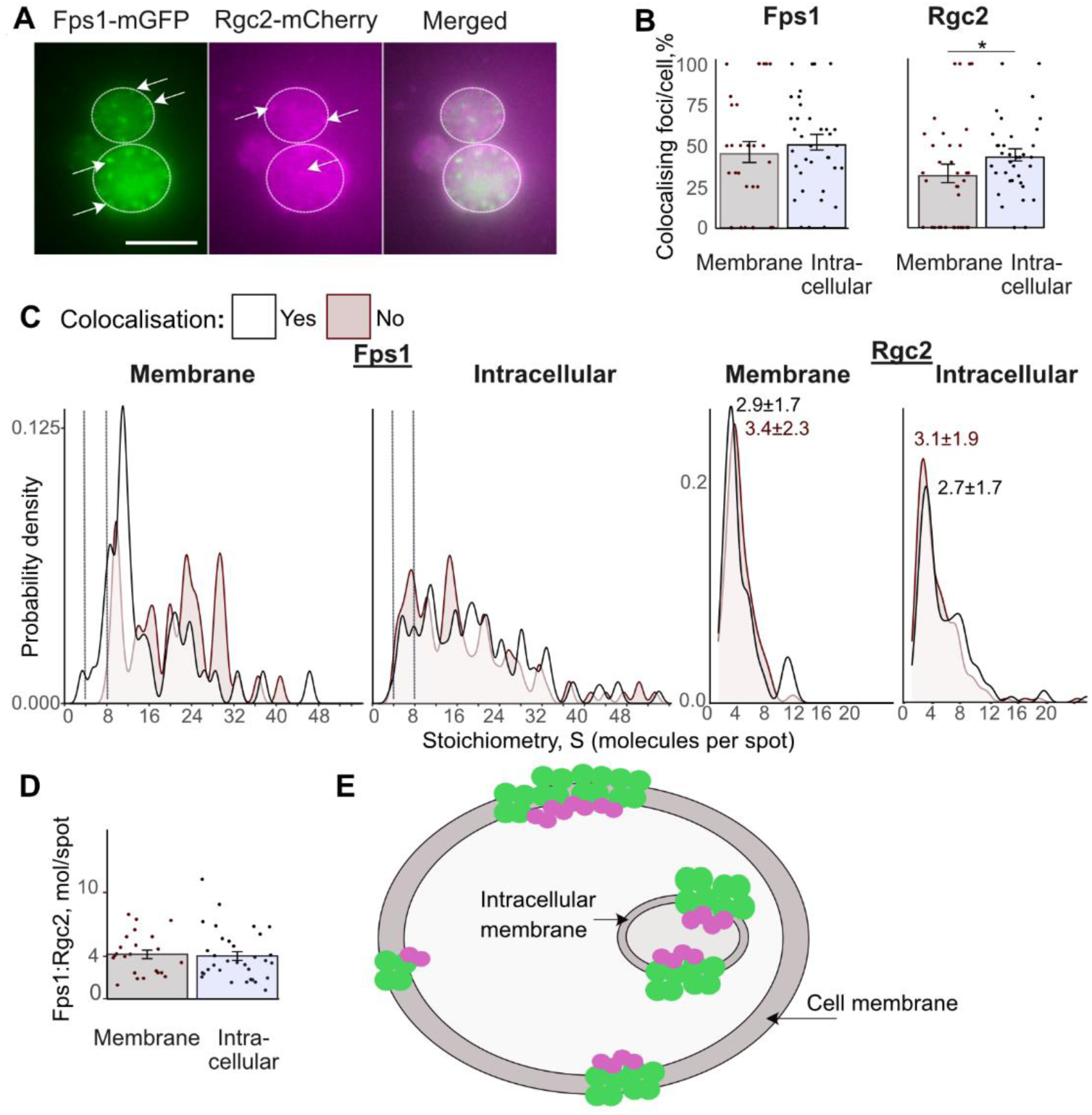
Fps1 and Rgc2 colocalise both on the membrane and inside the cell. **A.** Example microphotograph of the Fps1-mGFP Rgc2-mCherry strain. Scale bar 5 µm. White circles indicate cell boundaries; arrows point at exemplar fluorescent foci. **B.** Left: percent of all identified Fps1 foci on the membrane and inside the cell which colocalises with Rgc2. Dashed lines indicate stoichiometries of 4 and 8 molecules per spot. Right: percent of all identified Rgc2 foci on the membrane and inside the cell that colocalise with Fps1. Each datapoint reproduces one cell. Error bars represent standard error of mean. Wilcoxon rank sum test, *p<0.05. **C.** Kernel density plots of Fps1 and Rgc2 stoichiometry distributions on the membrane and inside the cell colocalised and non-colocalised with each other. Numbers indicate the highest peak value ±SE for Rgc2 foci colocalised (black) and non-colocalised (red) with Fps1. **D.** Molecular ratio of Fps1:Rgc2 within colocalised foci on the cell membrane and inside the cell. Error bars represent standard error of mean. **E.** Schematic representation of the Fps1-Rgc2 clusters identified on the cell membrane and intracellularly.

Fps1 and Rgc2 foci were found to colocalise with a high heterogeneity between cells, with 100% of Fps1 and Rgc2 foci colocalising in some cells and 0% foci colocalising in others (Fig. 2B). Overall, around 50% of the Fps1 foci were found to colocalise with Rgc2 foci regardless of the Fps1 localisation, while a higher proportion of Rgc2 foci were found colocalised with Fps1 in intracellular space to those found on the cell membrane (43% and 32%, respectively) (Fig. 2B). This is consistent with overall larger portion of Rgc2 located inside the cell (Supplementary Fig. 2A).

By quantifying the step-wise photobleaching of fluorophores (40), we estimated the number of Fps1 and Rgc2 molecules per identified focus (Fig. 2C and Supplementary Fig. 2B). Consistent with ours and others previous results (30, 41), our data indicates that Fps1 commonly consists of primarily tetrameric complexes with more distinct complex sizes at the membrane, while intracellular foci are also present as lower stoichiometry oligomers (Fig. 2C and Supplementary Fig. 2B). We applied the same technique to Rgc2-mCherry, revealing that Rgc2 mainly exists as a dimer, with apparent higher stoichiometry foci representing foci overlapping with the diffraction limit of the microscope or complexes with between 2 to 8 Rgc2 molecules (Fig. 2C and Supplementary Fig. 2B). This supports previous indications that Rgc2 may be organised into a dimer in order to activate Fps1 (16). Stoichiometries were similar regardless of colocalisation between Fps1 and Rgc2 (Fig. 2C).

We found that overall colocalisation ratio of Fps1:Rgc2 is 4.3±0.4 (±SE) on the membrane and 4.5±0.4 inside the cell (Fig. 2D, Table 1). This also suggests that the same Rgc2 dimer may act on two Fps1 tetramer clusters simultaneously.

**Table 1.** **Mean apparent stoichiometry of Fps1 and Rgc2 within colocalising foci** detected on the cellular membrane (membrane) and inside the cell (intracellular) in the wild type (wt) and *rgc1Δ* strains under standard (glucose) and oxidative stress (0.4 mM H_2_O_2_) conditions.

| Strain | Protein | Mean apparent stoichiometry, molecules/focus ± SE | | | |
| --- | --- | --- | --- | --- | --- |
|  |  | Glucose |  | H<sub>2</sub>O<sub>2</sub> |  |
|  |  | membrane | intracellular | membrane | intracellular |
| wt | Fps1 | 15.9±1.5 | 19.2±1.0 | 12.3±0.9 | 12.3±0.7 |
| | Rgc2 | 4.3±0.5 | 5.8±0.4 | 8.6±1.4 | 5.9±0.4 |
| *rgc1Δ* | Fps1 | 17.5±0.9 | 13.1±1.1 | 16.5±1.5 | 15.0±0.8 |
| | Rgc2 | 5.4±0.6 | 3.8±0.2 | 5.9±0.4 | 6.0±0.3 |

Taken together, under normal conditions, membrane architecture of the Fps1-Rgc2 complexes is represented by Fps1 (multi)tetramers and associated Rgc2 dimers (Fig. 2E). The presence of intracellular Fps1-Rgc2 clusters supports our previous idea of Fps1 residing on intracellular membranes possibly mediating substrate flux on organelles containing glycerol metabolising enzymes.

### Rgc1 regulates Fps1-Rgc2 localisation and intracellular behaviour

Due to the similarity of Rgc1 and Rgc2, it can be challenging to interpret mechanisms involving *RGC* genes simultaneously as they provide functional redundancy for each other, and phenotypes can thus not be linked to a separate gene. Hence, typically work on Fps1 regulation has utilised *rgc1Δ* strains (14–16, 20, 21, 28). To provide further insights into the organisation of Rgc proteins within the Fps1 gating system, we introduced the *RGC1* deletion into the Fps1-mGFP Rgc2-mCherry dual-coloured strain (Fig. 3A, Table 2). Surprisingly, the *RGC1* deletion had very limited effect on Rgc2 foci stoichiometry distribution (Fig. 3B), with Rgc2 still mainly present as a dimer both inside the cell and in association with the membrane. It has previously been reported that Rgc2 forms heterodimers with Rgc1 (16). Our data show no difference in the Fps1:Rgc2 stoichiometry ratios within colocalised foci between both the wildtype and *rgc1Δ* strains (Supplementary Fig. 3B). As the change in the peak membrane stoichiometry of Rgc2 is insignificant, hence, if Rgc1-Rgc2 heterodimers exist, they are present to a minor extent and overall, Rgc2 predominantly occurs as homodimers.

**Figure 3.**
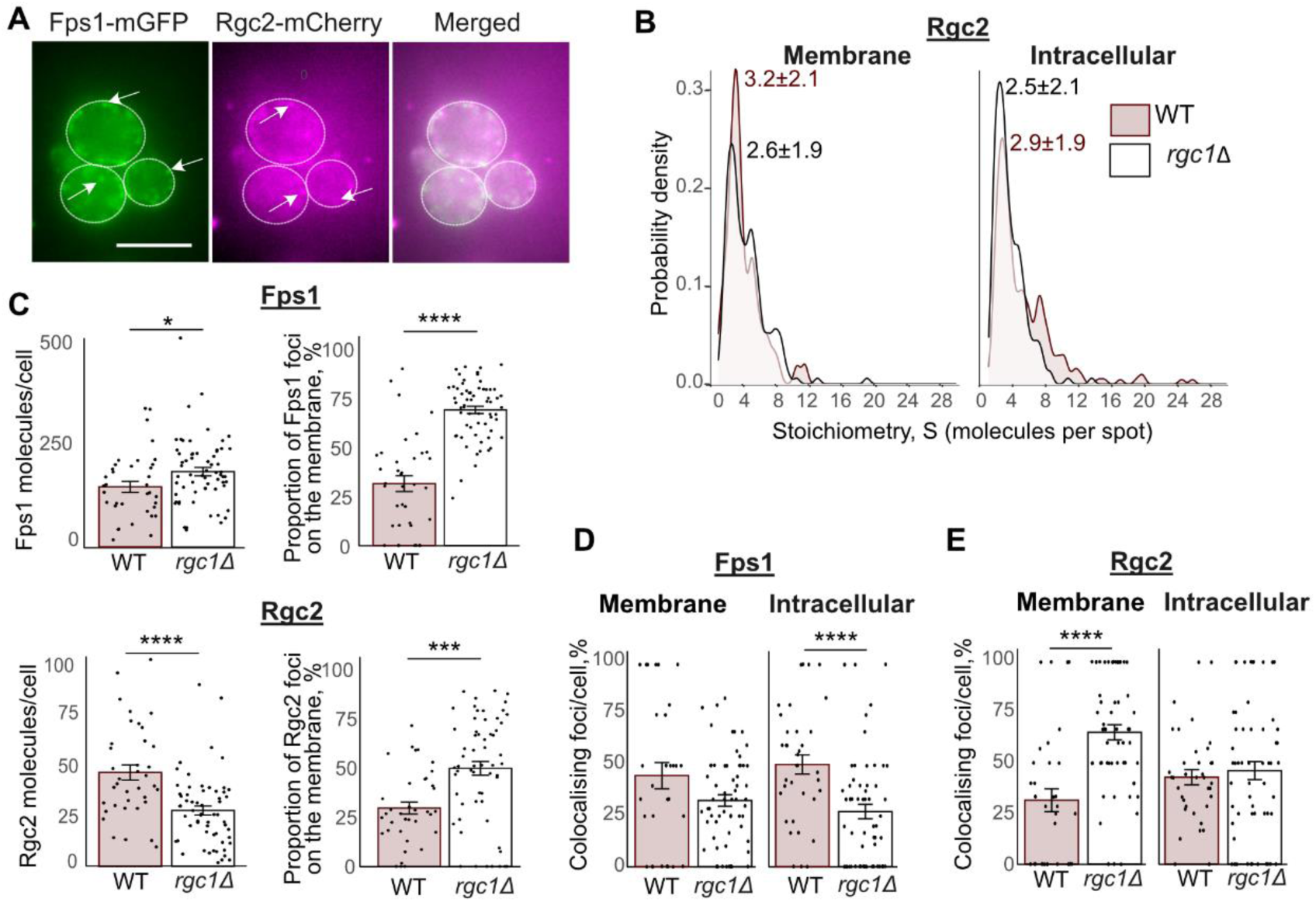
The effect of the RGC1 deletion on localisation and stoichiometry characteristics of the Fps1-Rgc2 clusters. **A.** Example microphotograph of the Fps1-mGFP Rgc2-mCherry *rgc1Δ* strain. Scale bar 5 µm. White circles indicate cell boundaries; arrows point at exemplar fluorescent foci. **B.** Kernel density plot of Fps1 and Rgc2 stoichiometry distributions on the membrane and inside the cell identified in the WT and *rgc1Δ* strains. Numbers indicate the highest peak value ±SE. **C.** Left: Total number of identified Fps1 (top) and Rgc2 (bottom) molecules per cell. Right: Fps1 (top) and Rgc2 (bottom) located on the membrane as a percent of all identified molecules per cell. Each datapoint reproduces one cell. Error bars represent standard error of mean. Wilcoxon rank sum test, *p<0.05, ***p≤0.001, ****p≤0.0001. **D.** Percent of all identified Fps1 foci on the membrane and (left) and intracellularly (right) in the wildtype (red) and the *rgc1Δ* strains that colocalise with Rgc2. Each datapoint reproduces one cell. Error bars represent standard error of mean. Wilcoxon rank sum test, ****p≤0.0001. **E.** Percent of all identified Rgc2 foci on the membrane and (left) and intracellularly (right) in the wildtype and the *rgc1Δ* strains that colocalise with Fps1. Each datapoint reproduces one cell. Error bars represent standard error of mean. Wilcoxon rank sum test, ****p≤0.0001.

**Table 2.** List of strains used in this study.

| Strain name | Genotype | Source |
| --- | --- | --- |
| Fps1-mGFP Rgc2-mCherry | *MATa BY4741 fps1::FPS1-mGFP HIS3 rgc2::RGC2-mCherry HYGROB* | This study |
| Fps1-mGFP Rgc2-mCherry *rgc1Δ* | *MATa BY4741 rgc1::KanMX fps1::FPS1-mGFP HIS3 rgc2::RGC2-mCherry HYGROB* | This study |
| Fps1-mGFP *rgc1Δ* | *MATa BY4741 rgc1::KanMX fps1::FPS1-mGFP HIS3* | This study |
| *rgc1Δ* | *MATa BY4741 rgc1::KanMX* | EUROSCARF(58) |

By summing up numbers of Fps1-mGFP molecules in all identified foci in a cell, we found that the *rgc1Δ* leads to an increase in Fps1 protein levels by ∼25%: 187±10 molecules per cell in the deletion strain as opposed to 149±14 molecules per cell in the wt (Fig. 3C, top left). Previous observations have indicated that the *rgc1Δ*/*rgc2Δ* double mutation results in Fps1 overexpression at 10-fold protein levels compared to the wildtype (15). This indicates that *rgc1Δ* produces an intermediate phenotype compared to the *rgc1Δ*/*rgc2Δ* double mutant. Conversely, the lack of Rgc1 causes fewer Rgc2 molecules that we were able to identify (Fig. 3C, bottom left), suggesting Rgc1 also regulated Rgc2 levels.

In the in *rgc1Δ* strain, we observed a clear shift in localisation of Fps1 and Rgc2 towards the cell membrane with fewer intracellular Fps1 and Rgc2 molecules (Fig. 3C, Supplementary Fig. 3C), suggesting that Rgc1 regulates Fps1/Rgc2 trafficking to the membrane. The *RGC1* deletion causes fewer colocalised Fps1 across the cell resulting in a higher number of free Fps1 (Fig. 3D and E). We also observed an increased proportion of colocalised Rgc2 in the membrane. Thereby, Rgc1 appears to regulate the Fps1-Rgc2 complex composition.

Taken together, Rgc1 regulates cellular localisation and spatial organisation of the Fps1-Rgc2 assemblies probably via controlling Rgc2 interactions with Fps1. Previous reports suggested that Rgc2 forms stable homodimers and possibly heterodimers with Rgc1. However, this conclusion was drawn based on the data obtained from overexpressed proteins in cell membrane only (16). We show that Rgc1 mainly affects the size of the Rgc2 clusters inside the cell. This suggests a potential role of Rgc1 in regulating the Fps1 gating system within the membranes of organelles containing glycerol metabolising enzymes. Moreover, our results indicate that it is predominantly Rgc2 homodimers that interact with Fps1 on the cell membrane.

### Intracellular Fps1-Rgc2 responses to H_2_O_2_ -induced oxidative stress in a Rgc1-dependent manner

Hydrogen peroxide, H_2_O_2_ is one of the common reactive oxygen species which when it accumulates in high enough concentrations, induces oxidative stress and following damages in eukaryotic cells (42, 43). H_2_O_2,_ has been shown to directly affect both Fps1 and Rgc2 (30, 44, 45). We previously have shown that oxidative stress causes Fps1 removal from the membrane and following degradation (30). Therefore, we sought to understand if the architecture of Fps1 and Rgc2 complexes changes upon cell exposure to H_2_O_2_ -induced oxidative stress (further denoted as “oxidative stress” (Fig. 4A)).

**Figure 4.**
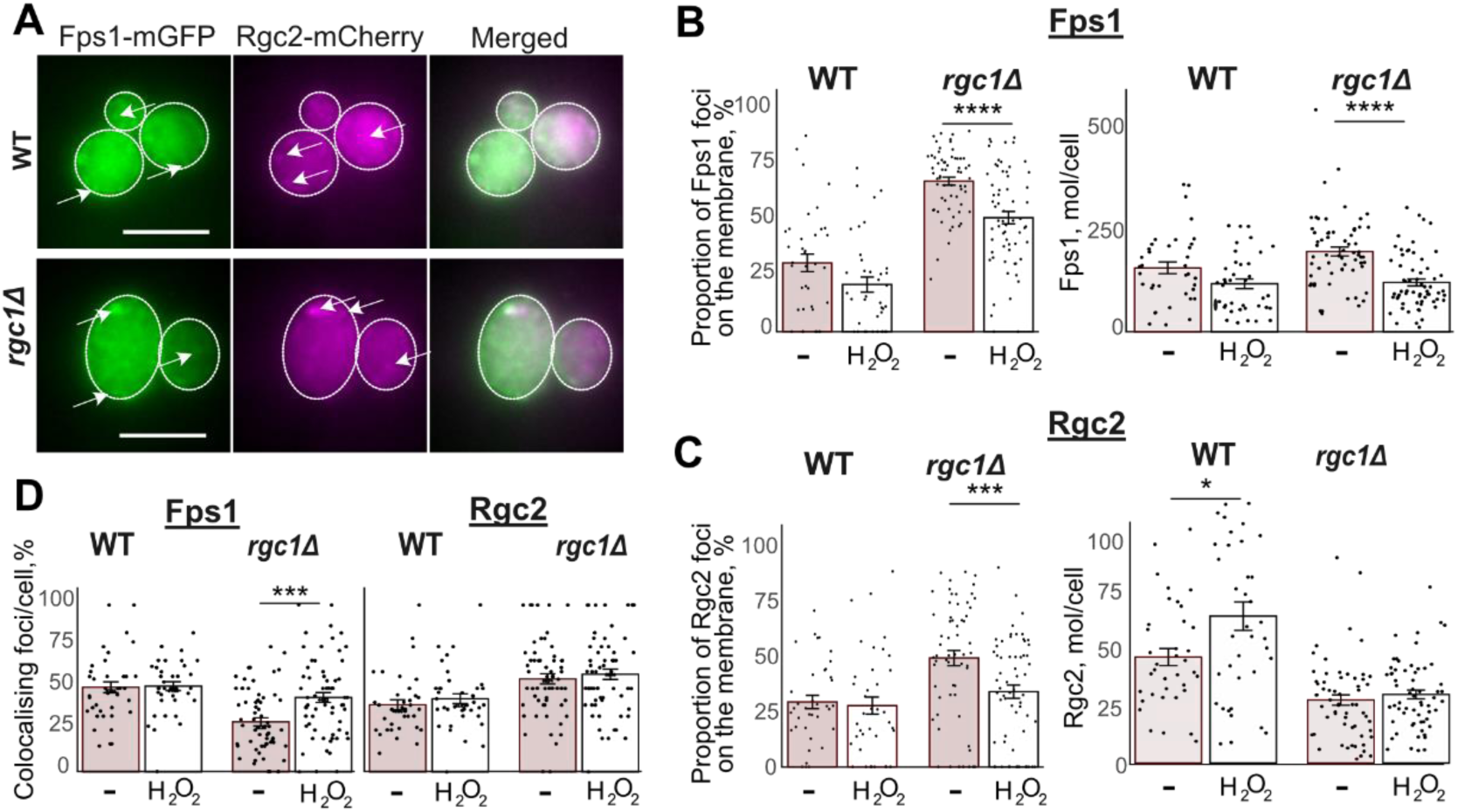
The effect of hydrogen peroxide on the Fps1-Rgc2 cluster organisation. **A.** Example microphotograph of the Fps1-mGFP Rgc2-mCherry wildtype (wt) and *rgc1Δ* strains upon exposure to 0.4 mM H_2_O_2_. Scale bar 5 µm. White circles indicate cell boundaries; arrows point at exemplar fluorescent foci. **B.** Left: Fps1 located on the membrane as a percent of all identified molecules per cell under normal conditions and 0.4 mM H_2_O_2_ exposure. Each datapoint reproduces one cell. Right: Total number of identified Fps1 molecules per cell under normal conditions and 0.4 mM H_2_O_2_ exposure. Error bars represent standard error of mean. Wilcoxon rank sum test, ****p≤0.0001. **C.** Left: Rgc2 located on the membrane as a percent of all identified molecules per cell under normal conditions and 0.4 mM H_2_O_2_ exposure. Each datapoint reproduces one cell. Right: Total number of identified Rgc2 molecules per cell under normal conditions and 0.4 mM H_2_O_2_ exposure. Error bars represent standard error of mean. Wilcoxon rank sum test, *p<0.05, ***p≤0.001. **D**. Left: percent of all identified Fps1 foci on the membrane and inside the cell which colocalises with Rgc2. Right: percent of all identified Rgc2 foci on the membrane and inside the cell that colocalise with Fps1. Each datapoint reproduces one cell. Error bars represent standard error of mean. Wilcoxon rank sum test, ***p≤0.001.

Consistent with our previous report, oxidative stress caused overall decrease in the mean apparent stoichiometry of Fps1(30) with significantly less Fps1 at the membrane, while the intracellular amount remained similar (Supplementary Fig. 4A and 4B). Rgc2 did not follow this behaviour with overall levels slightly increasing upon hydrogen peroxide exposure (Fig. 4C, Supplementary Fig. 4B). These changes, however, resulted in modest differences to stoichiometry distributions (Supplementary Fig. 4A).

Overall, upon oxidative stress, we were able to detect less Fps1 molecules, particularly at the membrane, (Fig. 4B, Supplementary Fig. 4B) regardless of the Rgc1 presence. In the wt, this has been previously linked to the internalisation and degradation of Fps1 (30). Hence Rgc1 is not essential in regulation of Fps1 localisation under oxidative stress.

We detected more Rgc2 under hydrogen peroxide exposure (Fig. 4C, right), mainly in the wildtype cells. Interestingly, upon Rgc1 absence, the increase in Rgc2 molecule numbers was only observed within the intracellular population (Supplementary Fig. 4B, bottom right), which also correlates with the decrease in the membrane portion of the Rgc2 foci (Fig. 4C, left). Hence, Rgc1 is involved in the Rgc2 response to H_2_O_2_-induced oxidative stress by affecting its’ localisation and Fps1-colocalisation profile. Oxidative stress does not affect the number of colocalised Fps1 or Rgc2 foci in wt background (Fig. 4D). However, in *rgc1Δ*, we identified more colocalised Fps1 and Rgc2 inside the cells (Supplementary Fig. 4C).

We show that upon *RGC1* absence, the overall levels of Rgc2 decrease, and those levels do not change upon H_2_O_2_ exposure. However, in the wt, oxidative stress increases the Rgc2 numbers. Hence, Rgc1 seems to promote Rgc2 stability in response to hydrogen peroxide stress.

## Discussion

Rgc1 and Rgc2 are known to be required to keep the Fps1 channel in an open state (15). Such open state allows the flux of known Fps1 substrates such as glycerol, acetic acid and toxic metalloids (6, 8–12). We show that the *rgc1Δ* mutation causes the total Fps1 numbers to increase along with a distinct shift in the Fps1 localisation towards the membrane (Fig. 3C, Supplementary Fig. 3C). Increased levels of Fps1 have been shown to increase its’ activity (11). At the same time, Rgc1 is also known to play a role in maintaining the Fps1 activity (15, 20, 21). Hence, the observed shift in Fps1 localisation in *rgc1Δ* coupled with the increase in total Fps1 could be compensatory for lowered Fps1 activity in the absence of Rgc1.

Surface protein turnover and levels are dependent on expression, post transcriptional modification, exocytosis, endocytosis and the recycling of proteins from early endosomes back onto the plasma membrane instead of being degraded (46–49). Thus, a decrease in Fps1 internalisation would lead to an accumulation of Fps1 at the membrane. Likewise, if Rgc1 normally acts to hinder Fps1 recycling back to the membrane that would also lead to more Fps1 in general and at the membrane specifically. Thereby the lowered numbers of intracellular Fps1, higher numbers of plasma membrane associated Fps1 and the increased Fps1 amount in *rgc1Δ* cells is not simply an increase in Fps1 production but due to a relative shift in Rgc1-dependent Fps1 internalization and recycling. Consistent with this observation, Fps1 protein levels have been shown to be stabilised from turnover upon cycloheximide treatment in an *RGC1/2* double deletion mutant (15).

Together these observations support the idea that Rgc1, and hence likely also its’ highly similar and functionally redundant paralog Rgc2, stabilises Fps1 levels, potentially by affecting Fps1 endocytosis and endosomal recycling. One mechanism which could explain the role of Rgc1 in Fps1activation and the observed phenotypes with a decreased endocytosis is that high cellular turgor is detrimental to endocytosis (50–53). Thus, a regulatory effect of Rgc1 on Fps1 endocytosis might simply be because the *rgc1Δ* mutant has a higher intracellular pressure due to decreased Fps1 activity.

Upon hydrogen peroxide exposure, Fps1 removal from the membrane and lower total levels have previously been shown to be due to Fps1 internalisation and degradation (30) Our data suggest that Fps1 is also removed from the membrane and degraded in a *rgc1Δ* mutant (Fig. 4B). Thus, the sole absence of Rgc1 cannot prevent Fps1 internalisation and degradation during oxidative stress.

When all Fps1 is mutated to a constitutively open mutant, *fps1-Δ1,* exposure to arsenic levels, survivable by wildtype cells, is lethal. (41) However, when co-expressed with *fps1-Δ1*, wildtype Fps1 is able to rescue these cells exposed to arsenic. We observed Fps1:Rgc2 are colocalised in foci of 4:1 (Fig. 2D, Table 1), implying that Rgc2 regulates multiple Fps1 at once. This would explain experiments with mixtures of wildtype and mutant Fps1, if in these cells, wildtype and mutant Fps1 form heterotetramers allowing Rgc2 to regulate mutant Fps1 through interactions with wildtype Fps1.

We also show that Rgc2 oligomerisation is primarily in the form of Rgc2 dimers and not Rgc1-Rgc2 heterodimers, complementing information from previous bulk assays (16). This also suggests a greater level of a separation between Rgc1 and Rgc2 functionality in terms of Fps1 regulation.

In summary, this study shows that dissecting the architecture and colocalisation characteristics of the key components if the Fps1-Rgc1/2 porin gating mechanism via single-molecule millisecond microscopy provides novel insights towards the underlying mechanisms of regulation. The role of Rgc1, and likely also Rgc2, is expanded towards a role in regulating Fps1 turnover under normal growth. As higher Fps1 levels in the pathogen *C. glabrata* are known to be directly involved in the pathogenic processes of acetate tolerance (23, 24) and antifungal uptake and efflux (54), the understanding of how Fps1 levels are regulated and maintained are of critical importance when designing antifungals targeting the Fps1-Rgc1/2 system. In addition, this study highlights importance of complementing genetic studies and bulk assays with *in vivo* single-molecule studies provide a better mechanistic understanding of a system.

## Methods

### Strain construction

The *rgc1Δ* deletion mutant expressing the Fps1-mGFP protein construct was designed in the same way as described previously (30, 55). We employed the monomeric version of GFP, mGFP, containing the A206K mutation which prevents potential self-oligomerisation of the fluorophore (56). To create the dual labelled strains, the mCherry::HygromycinB fragment flanked with 50bp up- and downstream the Rgc2 STOP codon sequences, was amplified from the pBS35 plasmid (Addgene plasmid #83797), purified and transformed into the Fps1-mGFP wt and *rgc1Δ* cells using homologous recombination (57). All strains used in this study are listed on Table 2.

### Growth conditions

Cells of the used strains from frozen stocks were pre grown overnight on Yeast Peptone Dextrose media (10 g/L Yeast Extract, 20 g/L Bacto Peptone, 2 % glucose (w/v), 20 g/L agar) at 30 °C. Cell maintenance was done on Yeast Nitrogen Base (YNB) medium (1× Difco™ YNB base, 5.0 g/L ammonium sulfate, pH 5.8-6.0) supplemented with 2 % glucose (w/v) and complete amino acid mixture (1× For-medium™ complete amino acid Supplement Mixture). For strain development, the amino acid mixture without histidine (1× For-medium™ amino acid -his drop-out Supplement Mixture) was used to select for Fps1-mGFP and 200 µg/mL Hygromycin B to select for Rgc2-mCherry.

For the microscopy experiments, the cells were first pre-cultured overnight in liquid YNB complete medium at 30 °C, 180 rpm, then grown to mid-logarithmic phase (OD_600_ 0.4-0.7) as described previously (30). Prior to imaging, cells were placed on a freshly cast pad of YNB media stabilised by 1 % agarose and immediately covered with a plasma-cleaned BK7 glass microscope coverslip (22 mm x 50 mm) and taken directly to the microscope. H_2_O_2_ stress was achieved by adding 0.4 mM H_2_O_2_ to the YNB media used to cast the agarose pad used for microscopy.

### Growth assay

Cells were grown overnight at 30°C at 250 rpm, in YNB-His medium supplemented with 4% glucose (w/v) with and without 1M sorbitol. Overnight cultures were diluted to OD_600_ 0.1 in respective media and grown until OD_600_ 0.3-0.5. Cells were then washed (1500g, 3min) and resuspend in YNB-His to OD_600_ 0.2. All strains were placed on YNB-His 2% glucose plates with and without 1M sorbitol, in five 10-fold dilutions 5µL spots. Plates were allowed to grow at 30°C and 37°C for 3 days.

### Single-molecule microscopy

We used a bespoke super-resolution fluorescence microscope, built around the body of a Nikon *Ti*-series epifluorescence microscope. Vortran 50 mW 473 nm and 561 nm wavelength lasers, coupled together using a dichroic mirror, were incident into the microscope with a 100x 1.49 NA Nikon objective lens to generate a collimated ∼20 µm (full width at half maximum) beam at the sample, with an intensity of typically 2.5–3 kW/cm^2^. The image was collected by a 300 mm focal length tube lens onto a Photometrics Evolve 512 Delta EMCCD camera, with a DV2 colour splitter to enable separate, simultaneous imaging of GFP and mCherry fluorescent protein components in the sample. The resulting magnification at the sample is 93 nm per pixel. The sample was held on a Mad City Labs XYZ positioning nanostage.

Sample preparation and super-resolved millisecond fluorescence microscopy was performed as previously published (30). To simultaneously visualise both channels, we utilised the Alternate Laser Excitation (ALEX) approach using 473 nm and 561 nm to excite mGFP and mCherry fluorophores, respectively (40, 59).The channel switching was done automatically with a 5ms cycle time between two channels and a resultant 10ms cycle time between each picture within a separate channel. Compared to (30) the analysis suite used updated parameters for foci detection and compartment segmentation. Any GFP spots found at the cell boundaries identified from the brightfield, we further refer to as “membrane”, while the spots found in the rest of the cell as “intracellular”. Foci stoichiometry was calculated using the principle of step-wise photobleaching of the fluorophores as described previously (30, 39).

### Data analysis

Foci were identified and tracked using a custom MATLAB program. In each frame, potential foci were detected through image transformation and thresholding via Otsu’s method. A 17x17 pixel region of interest (ROI) was established around each candidate to define the local background, while a central 5-pixel radius circle (the foreground) was processed with iterative Gaussian masking.

Foci were accepted if their signal-to-noise ratio was greater than 0.4, which was defined as the mean intensity of the foreground divided by the standard deviation of the background. Foci in subsequent frames were linked into trajectories based on the nearest neighbor, provided the distance was less than 5 pixels. These linked trajectories were considered "tracks" if they persisted for a minimum of four consecutive frames.

The characteristic intensity of mGFP or mCherry was established from the intensity values of foci near the end of photobleaching. This was confirmed by tracking foci beyond their disappearance and applying an edge-preserving filter to the raw intensity data to resolve the individual intensity steps caused by photobleaching. This value was then used to find the stoichiometry of foci by fitting a straight line to the first three intensity points and dividing the intercept by the characteristic intensity.

The GFP and mCherry images were aligned using the peak of the 2D cross-correlation between their corresponding brightfield images. Colocalisation between foci was determined as described previously (60).

Cells were segmented using a combination of edge detection of the brightfield image combined with watershedding (61).

Trajectory data was subsequently analysed using R version 4.2.1 (2022-06-23 ucrt). As a track can show up in two different cells if cells are overlapping the dataset was screened for duplicate tracks and a randomly selected duplicate of the tracks were excluded from analysis. As much of the data was predicted to not follow normal distribution, Wilcoxon signed ranks were used to evaluate statistics. For Kernel density estimation plots, the kernel width was set to 0.7 for both Fps1 and Rgc2. Colocalised tracks were defined using the overlap integral between fluorescent foci in both channels (62).

## Acknowledgements

Open access funding provided by the University of Gothenburg. This work was supported by Adlerbertska forskningsstiftelsen, the Biological Physical Sciences Interdisciplinary network (BPSI), the Royal Society Newton International Fellowship (grant number NF160208), Swedish research council (VR 2021-05201). MCL was supported by the Engineering and Physical Sciences Research Council EPSRC (EP/Y000501/1).

We would like to acknowledge the late Prof. Stefan Hohmann who first set us on the path of investigating Fps1 and Rgc2 interactions via the Slimfield super resolution microscopy.

MA: Conceptualization; Formal analysis; Funding acquisition; Investigation; Validation; Visualization; Writing - original draft. LV: Investigation, Visualization. AMA: Investigation. ML: Resources. AW: Software; Methodology; Formal analysis; Data curation; Supervision; Writing - review & editing SS: Methodology; Validation; Formal analysis; Data curation; Funding acquisition; Supervision; Writing - review & editing.

## Data availability

The data have been deposited in Zenodo (10.5281/zenodo.17423001) which includes growth assays and outputs of stoichiometry data analysis.

## Supplementary

**Supplementary Figure 1.**
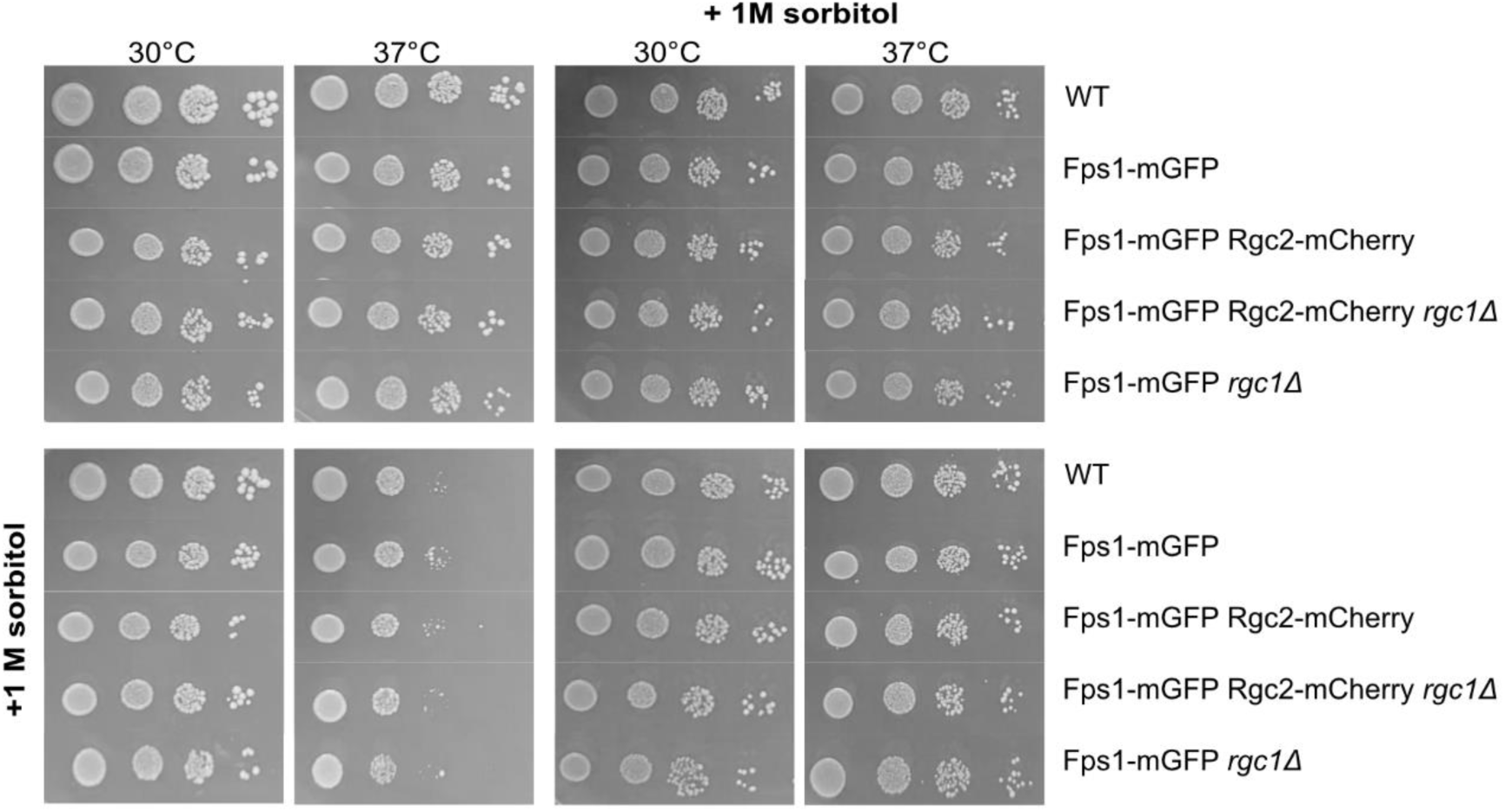
Characterisation of the strains developed for this study. Growth assessment of strains used in this study at 30 °C and 37°C on a standard (left) and sorbitol-supplemented (right) media. Prior to plating, cells were grown overnight with (bottom) and without (top) sorbitol addition. Each row represents series of 10-fold dilutions starting from OD_600_ 0.2.

**Supplementary Figure 2.**
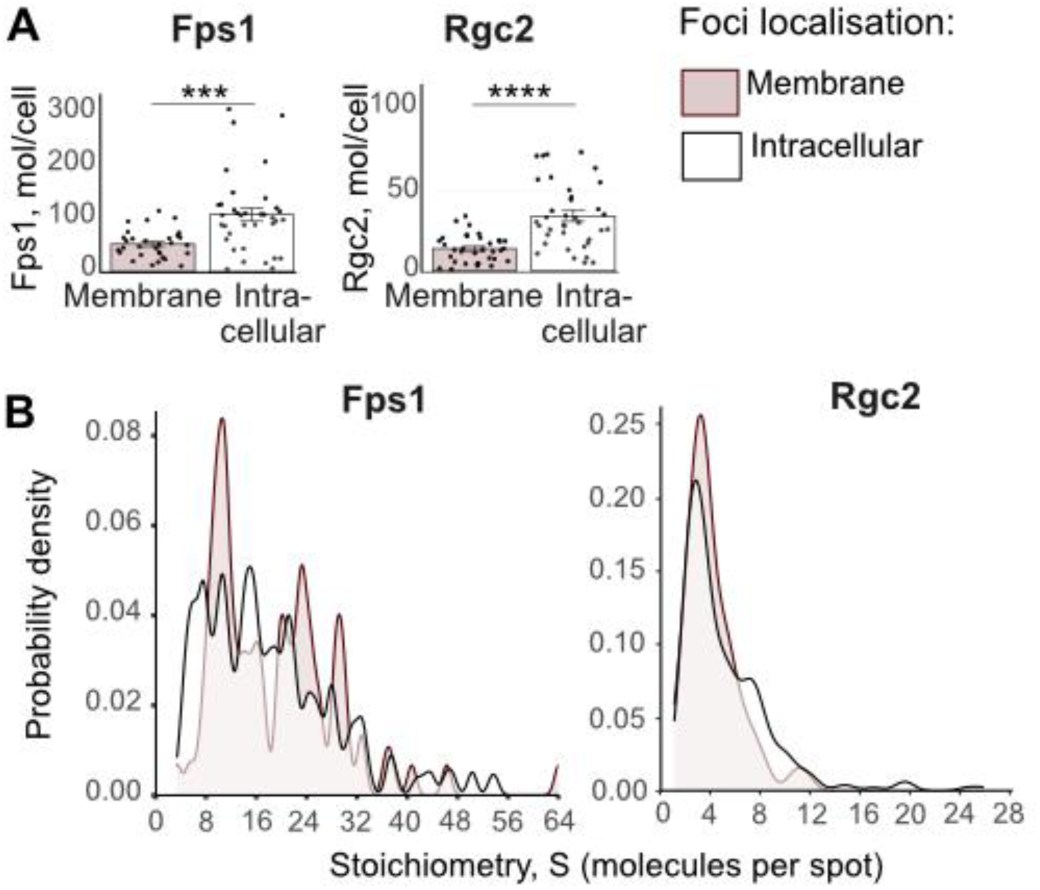
**A.** Total number of Fps1 (left) and Rgc2 (right) molecules within foci identified on the membrane or inside the cell. Error bars represent standard error of mean. Wilcoxon rank sum test, ***p≤0.001, ****p≤0.0001. **B.** Kernel density plots of the Fps1 (left) and Rgc2 (right) stoichiometry foci distributions detected on the cell membrane and inside the cell.

**Supplementary Figure 3.**
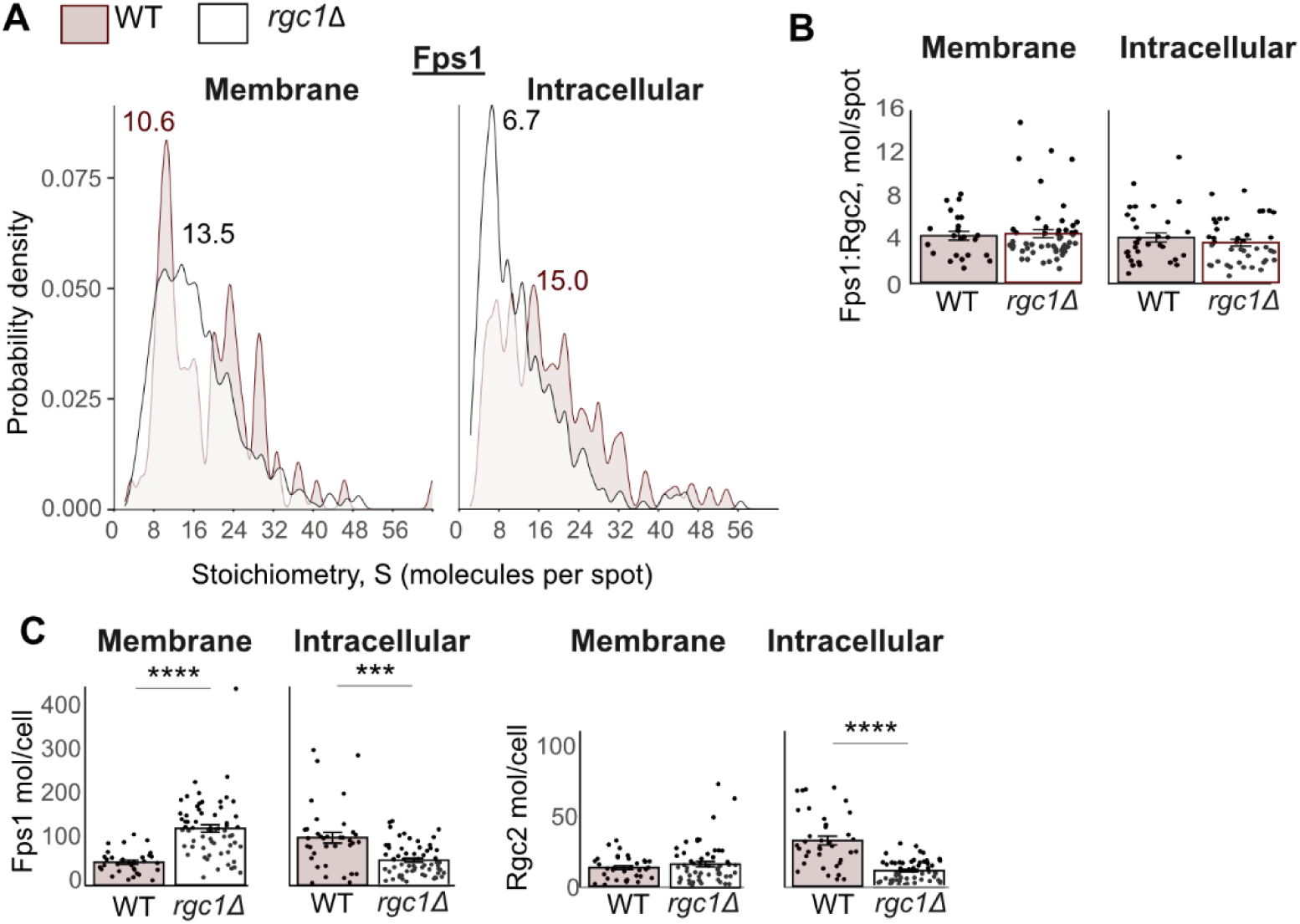
**A.** Kernel density plot of Fps1 stoichiometry distribution on the membrane and inside the cell identified in the WT and *rgc1Δ* strains. **B.** Molecular ratio of Fps1:Rgc2 within colocalised foci on the cell membrane and intracellularly in the wt and *rgc1Δ* strains. Error bars represent standard error of mean. **C.** Total number of identified Fps1 (left) and Rgc2 (right) molecules per cell on the cell membrane and intracellularly. Error bars represent standard error of mean. Wilcoxon rank sum test, ***p≤0.001, ****p≤0.0001.

**Supplementary Figure 4.**
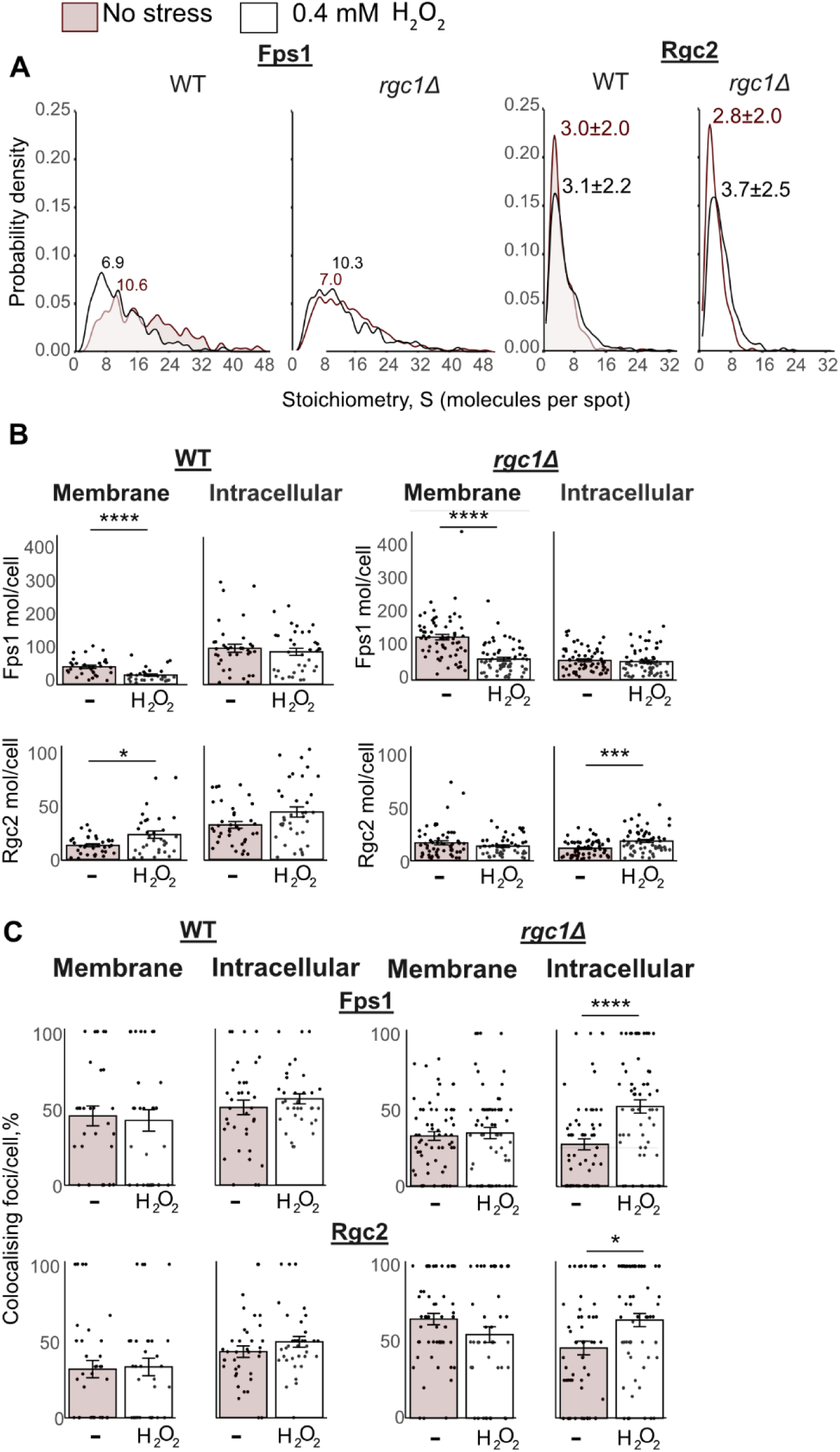
**A.** Kernel density plot of Fps1 and Rgc2 stoichiometry distributions in the WT and *rgc1Δ* strains under normal and 0.4mM H_2_O_2_ exposure conditions. **B**. The number of identified Fps1 (top) and Rgc2 (bottom) molecules per cell under normal conditions and 0.4 mM H_2_O_2_ exposure identified on the membrane and inside the cell in the WT (left) and *rgc1Δ* (right) strains. Error bars represent standard error of mean. Wilcoxon rank sum test, *p<0.05, ***p≤0.001, ****p≤0.0001. **C**. Top: percent of all identified Fps1 foci on the membrane and inside the cell which colocalises with Rgc2. Bottom: percent of all identified Rgc2 foci on the membrane and inside the cell that colocalise with Fps1. The data shown for the WT (left) and *rgc1Δ* (right) strains. Error bars represent standard error of mean. Wilcoxon rank sum test, *p<0.05, ****p≤0.0001.

**Supplementary Table 1.**
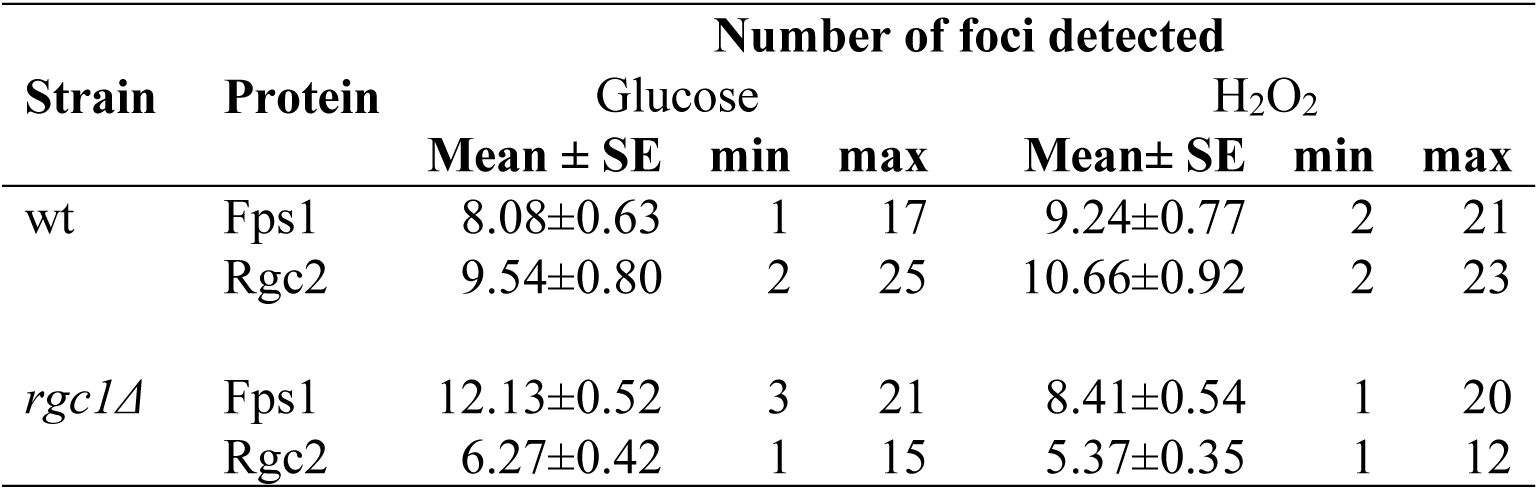
**Number of Fps1 and Rgc2 foci detected** in the wt and *rgc1Δ* strains under standard (glucose) and oxidative stress (0.4 mM H_2_O_2_) conditions. Mean as well as minimal (min) and maximal (max) numbers of identified foci are indicated.

## References

1. Ishibashi K, Morishita Y, Tanaka Y. 2017. The Evolutionary Aspects of Aquaporin Family, p. 35–50. In Yang, B (ed.), Aquaporins. Springer Netherlands, Dordrecht.

2. Agre P, Bonhivers M, Borgnia MJ. 1998. The Aquaporins, Blueprints for Cellular Plumbing Systems *. Journal of Biological Chemistry 273:14659–14662.

3. Zardoya R. 2005. Phylogeny and evolution of the major intrinsic protein family. Biol Cell 97:397–414.

4. Verkman AS. 2012. Aquaporins in Clinical Medicine. Annu Rev Med 63:303–316.

5. Pimpão C, Wragg D, da Silva I V, Casini A, Soveral G. 2022. Aquaglyceroporin Modulators as Emergent Pharmacological Molecules for Human Diseases. Front Mol Biosci Volume 9-2022.

6. Luyten K, Albertyn J, Skibbe WF, Prior BA, Ramos J, Thevelein JM, Hohmann S. 1995. Fps1, a yeast member of the MIP family of channel proteins, is a facilitator for glycerol uptake and efflux and is inactive under osmotic stress. EMBO J 14:1360–1371.

7. Mukhopadhyay R, Bhattacharjee H, Rosen BP. 2014. Aquaglyceroporins: Generalized metalloid channels. Biochim Biophys Acta Gen Subj 1840:1583–1591.

8. Wysocki R, Chéry CC, Wawrzycka D, Van Hulle M, Cornelis R, Thevelein JM, Tamás MJ. 2001. The glycerol channel Fps1p mediates the uptake of arsenite and antimonite in Saccharomyces cerevisiae. Mol Microbiol 40:1391–1401.

9. Mollapour M, Piper PW. 2007. Hog1 mitogen-activated protein kinase phosphorylation targets the yeast Fps1 aquaglyceroporin for endocytosis, thereby rendering cells resistant to acetic acid. Mol Cell Biol 27:6446–56.

10. Tamás MJ, Luyten K, Sutherland FC, Hernandez A, Albertyn J, Valadi H, Li H, Prior B a, Kilian SG, Ramos J, Gustafsson L, Thevelein JM, Hohmann S. 1999. Fps1p controls the accumulation and release of the compatible solute glycerol in yeast osmoregulation. Mol Microbiol 31:1087–104.

11. Maciaszczyk-Dziubinska E, Migdal I, Migocka M, Bocer T, Wysocki R. 2010. The yeast aquaglyceroporin Fps1p is a bidirectional arsenite channel. FEBS Lett 584:726–732.

12. Mollapour M, Shepherd A, Piper PW. 2009. Presence of the Fps1p aquaglyceroporin channel is essential for Hog1p activation, but suppresses Slt2(Mpk1)p activation, with acetic acid stress of yeast. Microbiology (N Y) 155:3304–3311.

13. Tsai M. 2014. Rgc1/Rgc2 deletions cause increased sensitivity to oxidative stress in saccaromyces cerevisiae, which can be overcome by constitutive nuclear Yap1 expression. ProQuest Dissertations and Theses. Boston University, Ann Arbor.

14. Lee J, Reiter W, Dohnal I, Gregori C, Beese-Sims S, Kuchler K, Ammerer G, Levin DE. 2013. MAPK Hog1 closes the S. cerevisiae glycerol channel Fps1 by phosphorylating and displacing its positive regulators. Genes Dev 27:2590–2601.

15. Beese SE, Negishi T, Levin DE. 2009. Identification of positive regulators of the yeast fps1 glycerol channel. PLoS Genet 5:e1000738.

16. Lee J, Levin DE. 2015. Rgc2 Regulator of Glycerol Channel Fps1 Functions as Homo- and Hetero-dimers with Rgc1. Eukaryot Cell 14:EC.00073-15.

17. Muir A, Roelants FM, Timmons G, Leskoske KL, Thorner J. 2015. Down-regulation of TORC2-Ypk1 signaling promotes MAPK-independent survival under hyperosmotic stress. Elife 4:1–13.

18. Abascal F, Ameliorate JL, Andersson M, Beitz E, Calamita G, Hohmann S, Nielsen S, Casini A, de Almeida A, De Ieso ML, Delporte C, Engel A, Frühbeck G, Huber VJ, Irisarri I, Kourghi M, Kwon T-H, Leonard A V., Madeira A, Marinelli RA, Méndez-Giménez L, Moon C, Moon D, Moura TF, Nyblom M, Pei J V., Rodríguez A, Rützler M, Tamma G, Törnroth-Horsefield S, Turner RJ, Vahedi-Faridi A, Valenti G, von Bülow J, Wacker S, Yool AJ, Zardoya R. 2018. Aquaporins in health and disease: new molecular targets for drug discovery, 1st ed. CRC Press, Boca Raton, FL.

19. Brewster JL, Gustin MC. 2014. Hog1: 20 years of discovery and impact. Sci Signal 7:re7–re7.

20. Baltanás R, Bush A, Couto A, Durrieu L, Hohmann S, Colman-Lerner A. 2013. Pheromone-induced morphogenesis improves osmoadaptation capacity by activating the HOG MAPK pathway. Sci Signal 6:1–30.

21. Dunayevich P, Baltanás R, Clemente JA, Couto A, Sapochnik D, Vasen G, Colman-Lerner A. 2018. Heat-stress triggers MAPK crosstalk to turn on the hyperosmotic response pathway. Sci Rep 8:15168.

22. Alastruey-Izquierdo A, Organization WH, Organization WH. 2022. WHO fungal priority pathogens list to guide research, development and public health action.

23. Mota S, Alves R, Carneiro C, Silva S, Brown AJ, Istel F, Kuchler K, Sampaio P, Casal M, Henriques M, Paiva S. 2015. Candida glabrata susceptibility to antifungals and phagocytosis is modulated by acetate. Front Microbiol Volume 6–2015.

24. Bernardo RT, Cunha D V, Wang C, Pereira L, Silva S, Salazar SB, Schröder MS, Okamoto M, Takahashi-Nakaguchi A, Chibana H, Aoyama T, Sá-Correia I, Azeredo J, Butler G, Mira NP. 2017. The CgHaa1-Regulon Mediates Response and Tolerance to Acetic Acid Stress in the Human Pathogen Candida glabrata. G3 Genes|Genomes|Genetics 7:1–18.

25. Sara E. B-S, Shih-Jung P, Jongmin L, Elizabeth H-W, P. CB, E. LD. 2012. Mutants in the Candida glabrata Glycerol Channels Are Sensitized to Cell Wall Stress. Eukaryot Cell 11:1512–1519.

26. Costa C, Ponte A, Pais P, Santos R, Cavalheiro M, Yaguchi T, Chibana H, Teixeira MC. 2015. New Mechanisms of Flucytosine Resistance in C. glabrata Unveiled by a Chemogenomics Analysis in S. cerevisiae. PLoS One 10:e0135110.

27. Lee J, Levin DE. 2022. Differential metabolism of arsenicals regulates Fps1-mediated arsenite transport. Journal of Cell Biology 221:e202109034.

28. Ahmadpour D, Maciaszczyk-Dziubinska E, Babazadeh R, Dahal S, Migocka M, Andersson M, Wysocki R, Tam??s MJ, Hohmann S. 2016. The mitogen-activated protein kinase Slt2 modulates arsenite transport through the aquaglyceroporin Fps1. FEBS Lett 590:3649–3659.

29. Shashkova S, Leake MC. 2018. Systems biophysics: Single-molecule optical proteomics in single living cells. Curr Opin Syst Biol 7.

30. Shashkova S, Andersson M, Hohmann S, Leake MC. 2020. Correlating single-molecule characteristics of the yeast aquaglyceroporin Fps1 with environmental perturbations directly in living cells. Methods 10.1016/j.ymeth.2020.05.003.

31. Plank M, Wadhams GH, Leake MC. 2009. Millisecond timescale slimfield imaging and automated quantification of single fluorescent protein molecules for use in probing complex biological processes. Integrative Biology 1:602–612.

32. Wollman AJM, Syeda AH, Howard JAL, Payne-Dwyer A, Leech A, Warecka D, Guy C, McGlynn P, Hawkins M, Leake MC. 2024. Tetrameric UvrD Helicase Is Located at the E. Coli Replisome due to Frequent Replication Blocks. J Mol Biol 436:168369.

33. Jin X, Lee J-E, Schaefer C, Luo X, Wollman AJM, Payne-Dwyer AL, Tian T, Zhang X, Chen X, Li Y, McLeish TCB, Leake MC, Bai F. 2025. Membraneless organelles formed by liquid-liquid phase separation increase bacterial fitness. Sci Adv 7:eabh2929.

34. Wollman AJM, Muchová K, Chromiková Z, Wilkinson AJ, Barák I, Leake MC. 2020. Single-molecule optical microscopy of protein dynamics and computational analysis of images to determine cell structure development in differentiating Bacillus subtilis. Comput Struct Biotechnol J 18:1474–1486.

35. Syeda AH, Wollman AJM, Hargreaves AL, Howard JAL, Brüning J-G, McGlynn P, Leake MC. 2019. Single-molecule live cell imaging of Rep reveals the dynamic interplay between an accessory replicative helicase and the replisome. Nucleic Acids Res 47:6287–6298.

36. Wollman AJM, Leake MC. 2022. Single-Molecule Narrow-Field Microscopy of Protein-DNA Binding Dynamics in Glucose Signal Transduction of Live Yeast Cells, p. 5–16. In Leake, MC (ed.), Chromosome Architecture: Methods and Protocols. Springer US, New York, NY.

37. Shashkova S, Nyström T, Leake MC, Wollman AJM. 2021. Correlative single-molecule fluorescence barcoding of gene regulation in Saccharomyces cerevisiae. Methods 193:62–67.

38. Wollman AJM, Hedlund EG, Shashkova S, Leake MC. 2020. Towards mapping the 3D genome through high speed single-molecule tracking of functional transcription factors in single living cells. Methods 170:82–89.

39. Wollman AJM, Shashkova S, Hedlund EG, Friemann R, Hohmann S, Leake MC. 2017. Transcription factor clusters regulate genes in eukaryotic cells. Elife 6:e27451.

40. Wollman AJM, Shashkova S, Welkenhuysen N, Hedlund EG, Hohmann S, C. Leake M. 2017. Time-Resolved Single Cell, Sub-Cellular Compartmentalized Proteomics, Combining Precise Microfluidics, Deconvolution and Ultrasensitive Single-Molecule Microscopy. Biophys J 112:313a.

41. Beese-Sims SE, Lee J, Levin DE. 2011. Yeast Fps1 glycerol facilitator functions as a homotetramer. Yeast 28:815–819.

42. Saputra F, Kishida M, Hu SY. 2024. Oxidative stress induced by hydrogen peroxide disrupts zebrafish visual development by altering apoptosis, antioxidant and estrogen related genes. Sci Rep 14.

43. Xin X, Gong T, Hong Y. 2022. Hydrogen peroxide initiates oxidative stress and proteomic alterations in meningothelial cells. Sci Rep 12.

44. Cohen TJ, Lee K, Rutkowski LH, Strich R. 2003. Ask10p mediates the oxidative stress-induced destruction of the Saccharomyces cerevisiae C-type cyclin Ume3p/Srb11p. Eukaryot Cell 2:962–970.

45. Jin C, Strich R, Cooper KF. 2014. Slt2p phosphorylation induces cyclin C nuclear-to-cytoplasmic translocation in response to oxidative stress. Mol Biol Cell 25:1396–1407.

46. Laidlaw KME, MacDonald C. 2018. Endosomal trafficking of yeast membrane proteins. Biochem Soc Trans 46:1551–1558.

47. Martin-Perez M, Villén J. 2017. Determinants and Regulation of Protein Turnover in Yeast. Cell Syst 5:283–294.e5.

48. Delic M, Valli M, Graf AB, Pfeffer M, Mattanovich D, Gasser B. 2013. The secretory pathway: exploring yeast diversity. FEMS Microbiol Rev 37:872–914.

49. Ma M, Burd CG. 2020. Retrograde trafficking and plasma membrane recycling pathways of the budding yeast Saccharomyces cerevisiae. Traffic 21:45–59.

50. Lemière J, Chang F. 2023. Quantifying turgor pressure in budding and fission yeasts based upon osmotic properties. Mol Biol Cell 34.

51. Carlsson AE, Bayly P V. 2014. Force generation by endocytic actin patches in budding yeast. Biophys J 106:1596–1606.

52. Aghamohammadzadeh S, Ayscough KR. 2009. Differential requirements for actin during yeast and mammalian endocytosis. Nat Cell Biol 11:1039–42.

53. Dmitrieff S, Nédélec F. 2015. Membrane Mechanics of Endocytosis in Cells with Turgor. PLoS Comput Biol 11.

54. Teixeira V, Costa V. 2015. Unraveling the role of the Target of Rapamycin signaling in sphingolipid metabolism. Prog Lipid Res 61:109–133.

55. Shashkova S, Nyström T, Leake MC. 2022. Copy Number Analysis of the Yeast Histone Deacetylase Complex Component Cti6 Directly in Living Cells. Methods in Molecular Biology 2476:183–190.

56. Zacharias DA, Violin JD, Newton AC, Tsien RY. 2002. Partitioning of lipid-modified monomeric GFPs into membrane microdomains of live cells. Science (1979) 296:913–916.

57. Gietz RD, Schiestl RH. 2007. Frozen competent yeast cells that can be transformed with high efficiency using the LiAc/SS carrier DNA/PEG method. Nature Protocols 2007 2:1 2:1–4.

58. Brachmann CB, Davies A, Cost GJ, Caputo E, Li J, Hieter P, Boeke JD. 1998. Designer deletion strains derived from Saccharomyces cerevisiae S288C: A useful set of strains and plasmids for PCR-mediated gene disruption and other applications. Yeast 14:115–132.

59. Leake MC, Chandler JH, Wadhams GH, Bai F, Berry RM, Armitage JP. 2006. Stoichiometry and turnover in single, functioning membrane protein complexes. Nature 443:355–358.

60. Wollman AJM, Fournier C, Llorente-Garcia I, Harriman O, Payne-Dwyer AL, Shashkova S, Zhou P, Liu T-C, Ouaret D, Wilding J, Kusumi A, Bodmer W, Leake MC. 2022. Critical roles for EGFR and EGFR–HER2 clusters in EGF binding of SW620 human carcinoma cells. J R Soc Interface 19:20220088.

61. Wollman AJM, Kioumourtzoglou D, Ward R, Gould GW, Bryant NJ. 2022. Large scale, single-cell FRET-based glucose uptake measurements within heterogeneous populations. iScience 25.

62. Haapasalo K, Wollman AJM, de Haas CJC, van Kessel KPM, van Strijp JAG, Leake MC. 2019. Staphylococcus aureus toxin LukSF dissociates from its membrane receptor target to enable renewed ligand sequestration. The FASEB Journal 33:3807–3824.

